# Antifungal activity of selected lactic acid bacteria from olive drupes

**DOI:** 10.1101/2022.11.07.515451

**Authors:** Mario Riolo, Carlos Luz, Elena Santilli, Giuseppe Meca, Santa Olga Cacciola

**Affiliations:** Department of Agriculture, Food and Environment, University of Catania, 95123 Catania, Italy; Department of Agricultural Science, Mediterranean University of Reggio Calabria, 89122 Reggio Calabria, Italy; Laboratory of Food Chemistry and Toxicology, Faculty of Pharmacy, University of Valencia, Burjassot, Spain; Council for Agricultural Research and Economics, Research Centre for Olive, Fruit and Citrus crops (CREA-OFA), 87036 Rende, Cosenza, Italy

**Keywords:** LABs, VOCs, Organic Compounds, Phenolic Acids, olive diseases

## Abstract

In this study, 16 Lactobacilli (LABs) isolated from the drupes of olive (*Olea europaea*) oil varieties were identified as *Lactiplantibacillus plantarum* (seven isolates), *Pediococcus pentosaceus* (six isolates), *Enterococcus faecium* (two isolates) and *Streptococcus salivarius* (a single isolate) by peptide mass fingerprinting and sequencing of the 16S rRNA. Antifungal activity of LABs and their cell-free fermentates (CFSs) against several plant pathogenic fungi and oomycetes (fungi *sensu lato*), including *Alternaria, Aspergillus Colletotrichum, Penicillium, Plenodomus* and *Phytophthora*, was evaluated *in vitro* using the culture overlaying and the agar diffusion tests. Minimal inhibitory concentration (MIC) and minimal fungicidal concentration (MFC) were determined. LABs showed antifungal activity against the fungi *sensu lato* tested. The most noticeable inhibitory activity was shown by isolates of *L. plantarum* and *P. pentosaceus* against *Fusarium oxysporum, Colletotrichum* species and *Penicillium nordicum*. Chemical analysis revealed CFSs contained acid lactic and variable quantities of 14 diverse phenolic acids and 26 volatile organic compounds (VOCs). No obvious correlation was found between the metabolic profile of LABs and their antifungal efficacy. However, it is the first time that the potential of fermentates of LABs, recovered from drupes of olive oil varieties, as natural fungicides, was demonstrated.

## 1. Introduction

Several fungi and oomycetes (fungi *sensu lato, s*.*l*.) cause diseases on olive (*Olea europaea* L., family *Oleaceae*) (Cacciola et al., 2012; Santilli et al., 2020; Scanu et al., 2021; Schena et al., 2011; Riolo et al., 2022). Some of these pathogens, such as *Alternaria, Aspergillus, Colletotrichum, Fusarium* and *Penicillium*, are responsible for decay of olive drupes and deterioration of oil quality (Angerosa et al., 1999; Baffi et al., 2012; Fakas et al., 2010; Markakis et al., 2010; F. Peres et al., 2021; Romero et al., 2022; Torbati et al., 2014). The damage caused by these fungi has a particular economic relevance in countries where high quality standards of olive oil are pursued, such as Spain and Italy, which with Greece are the three major olive producer countries worldwide (Fraga et al., 2021). Presently, the management of olive diseases caused by fungi *s*.*l*., including fruit rots, largely relies on pre-harvest treatments with copper and synthetic fungicides. However, European Directive 2009/128/EC, aiming at reducing substantially the use of pesticides in agriculture, has fostered the search of alternative, environmentally friendly and toxicologically more safe strategies, such as the use of generally recognized as safe (GRAS) natural substances and microorganisms or their metabolites inciting natural plant defence mechanisms (La Spada et al., 2021; Lin et al., 2021; López-Moral et al., 2021; López-Velázquez et al., 2021; Moral et al., 2018; Pakora et al., 2018; Pangallo et al., 2017; Ren et al., 2022; Stracquadanio et al., 2020). Lactobacilli (LABs) is the term commonly adopted to refer to Gram-positive bacteria formerly grouped in the genus *Lactobacillus* (family, *Lactobacillaceae*), the largest genus of lactic acid bacteria that recently was split into several genera, according to new molecular taxonomic criteria (Zheng et al., 2020). LABs are found in a wide variety of environments, including soil (most associated with the rhizosphere), plants (particularly decaying plant material), animals and humans (especially the oral cavity, intestinal tract, and vagina). LABs are widely applied in food processing as starter cultures of fermentations and additives, and in fruit and vegetable supply chains have been exploited as food preservative for their ability to prevent the microbe dependent food spoilage (Liu et al., 2017). The efficacy of LABs in prolonging the fruit shelf-life depends on their ability to produce antibacterial, antioxidant and antifungal secondary metabolites. LABs are also used to improve the flavour and taste and as probiotics to enhance the nutritional value of foods (Dopazo et al., 2021; Erten et al., 2014; Qi et al., 2021). Previous studies highlighted the diversity of populations of LABs in the microbioma of olive fruit and showed it was dependent on the olive cultivar and geographical origin (Hurtado et al., 2008, 2012; C. M. Peres et al., 2014). Predominant species of LABs in most olive fermentations are *Lactiplantibacillus plantarum* and *Pediococcus pentosaceus* (Aponte et al., 2012; Ciafardini et al., 1994; Heperkan, 2013; Hurtado et al., 2012; Lanza, 2013; Leal-Sánchez et al., 2003; Marsilio et al., 2005; Medina et al., 2008; Ruiz-Barba et al., 2010). LABs comprise numerous GRAS species including the latter ones (Behera et al., 2018; Jiang et al., 2021).

The objectives of this study were (i) to identify LABs isolated from drupes of oil varieties of olive, (ii) evaluate the *in vitro* inhibitory activity of these LABs and their fermentates against fungi *s*.*l*. pathogens of olive and other crop plants, (iii) assess the ability of these LABs to produce bioactive metabolites, such as volatile organic compounds (VOCs), organic and phenolic acids.

## 2. Materials and Methods

### 2.1 Bacterial and fungal strains

LABs characterised in this study were obtained from different olive varieties (var) or cultivars (cv) sourced in Italy and Spain. The isolation was performed using sterile urine baker containing olives and Man Rogosa and Sharpe broth (MRS-B) (Oxoid, Dublin, Ireland) (ratio 1:2). Three replicates were performed for each olive sample. They were incubated at 37°C for 72 h in an anaerobic atmosphere, using an Anaerocult® system (Millipore, Milan, Italy). Serial dilutions (10^−4^, 10^−5^, 10^−6^) of the fermented media were sprayed on Man Rogosa and Sharpe agar (MRS-A) in Petri dishes and kept as before for 48 h. For each plate, five colonies were isolated and grown individually on MRS-A (Oxoid, Dublin, Ireland) in the same incubation conditions. To obtain pure cultures, different colony forming units were sub-cultured on MRS-A. Gram stain was performed on isolated cultures to select only Gram-positive bacteria (Forbes et al., 1998). Gram-positive, catalase-negative bacilli were identified as putative LABs. Selected cultures were preserved in cryotubes on MRS-B submerged with 30% glycerol at −20°C.

To restore the cultures, frozen bacteria were added to MRS-B and incubated for 24 h at 37°C, then 1 mL aliquots were transferred to MRS-B and incubated as above.

To evaluate the antifungal activity of isolates, LABs were tested *in vitro* against pathogenic fungi *s*.*l*., i.e. *Alternaria alternata* (isolates CECT 646 and ITEM 8121), *Aspergillus flavus* (ITEM 8111), *Colletotrichum acutatum* (UWS149 and C9D2C), *C. fiorinae* (ER2147 and C15D6A), *C. gloeosporioides* (C2 and RD9B), *C. nymphaeae* (RB 012 and RB 428), *Fusarium oxysporum* (COAL 68), *Penicillium digitatum* (P1PP0 and N2F1), *P. expansum* (COAL 95), *P. nordicum* (CECT 2320), *Plenodomus tracheiphilus* (Pt2), *Phytophthora nicotianae* (P4K3F8 and T-2K1A) and *Ph. oleae* (V-2K10A). Those isolates were sourced from the collection of the Molecular Plant Pathology laboratory of the Department of Agriculture, Food and Environment, Catania University, Italy, the Colección Española de Cultivos Tipo (CECT), Valencia University, Spain, and ITEM Collection from Istituto di Scienze delle Produzioni Alimentari (ISPA), National Research Council, Bari, Italy. Most of them were characterized in previous studies (El boumlasy et al., 2022; La Spada et al., 2021; Carlos Luz et al., 2021; Migheli et al., 2009; Riolo et al., 2020, 2021, 2022).

### 2.2 Bacterial identification

LAB strains characterization was performed by extraction of bacterial cultures, as reported by Maier et al. (2006). The method was performed with MALDI-TOF MS using a Microflex L20 mass spectrophotometer (Bruker Daltonics, Billerica, MA, USA) equipped with an N2 laser. Spectra were acquired in positive linear ion mode. Voltage acceleration was 20 kV and the mass range for the analysis was delimited from 2000 to 20000 Da. For each sample, three spectra were obtained following the method MALDI Biotyper Realtime Classification (RTC). Identification corresponded to the largest log score. Results were compared with the database MBT 7854 y MBT 7311_RUO (Bruker Daltonics).

### 2.3 Fungal inoculum

Ascomycota isolates were grown on potato dextrose agar (PDA; Oxoid Ltd., Basingstoke, UK) at 25°C in the dark until the mycelium covered the Petri dish. A conidial suspension (10^4^ ml^−1^) of each isolate in sterile distilled water (SDW), used as inoculum, was determined with a Neubauer chamber. *Phytophthora nicotianae* and *Ph. oleae* were grown on Oat Meal Agar (OMA, Thermo Fisher, Waltham, Massachusetts, USA) at 22°C in the dark for 7 days. Zoospore production was performed according to Aloi et al. (2021). Zoospore concentration was determined with a Neubauer chamber and adjusted to 10^4^ zoospores/mL.

### 2.4 Preparation of CFS (culture-free supernatants)

The preparation of culture-free supernatants (CFSs) was performed according Dopazo et al., (2021). Then, the medium was centrifuged at 4°C and 4000 rpm for 15 min, in an Eppendorf 5810 R centrifuge (Eppendorf, Hamburg, Germany). CFS was recovered, frozen at −80°C, lyophilized in a FreeZone 2.5 L Benchtop Freeze Dry System, (Labconco, Kansas City, MO, USA) and preserved at −20 °C for further analysis. To be tested lyophilized CFSs were resuspended in SDW.

### 2.5 Qualitative assay of antifungal activity

The effect of LABs and their CFSs on the growth of tested fungi *s*.*l*. was preliminarily evaluated using two distinct qualitative methos, the culture overlay, and the diffusion agar tests, respectively. In the first test, LAB cultures, grown for 24 h at 37ºC, were inoculated in the centre of MRS-A dishes containing 15 ml of medium and incubated at 37ºC for 72 h. Conidia or zoospores were then suspended in sterile 0.1% Tween–water and their concentration was adjusted using a Neubauer chamber. After 24 h incubation, dishes inoculated with the LAB suspension were covered with 15 mL of PDA (45ºC) containing 10^4^ spores/mL and incubated at 25ºC for another 72 h. After incubation the fungal growth inhibition was determined according to Guimarães et al. (2018). The diffusion agar test was used to assay the inhibitory effect of bacterial CFSs on fungi *s*.*l*. tested according to Dopazo et al. (2021). Halos larger than 8 mm were considered positive for antifungal activity (Varsha et al., 2014). The results of the test, performed in triplicate, were scored based on an empirical scale where: (+) mean diameter of inhibition halo <8 mm; (++) mean diameter of inhibition halo 8-10 mm; (+++) mean diameter of inhibition halo >10 mm.

### 2.6 Quantitative assay of antifungal activity

The minimal inhibitory concentration (MIC) and minimal fungicidal concentration (MFC) of the CFSs were determined in sterile 96-well microplates using the method described by Luz et al. (2020). For each well, 100 μL of CSF were dispensed at doses between 0.1 and 200 g/L. Then, 100 μL of conidial or zoospore suspension (5 × 10^4^ spores/mL) in PDB were added to each well. A negative control was obtained by adding 200 μL PDB to a well and a positive one by adding 100 μL PDB and 100 μL of the spore suspension. Microplates were incubated at 25°C for 72 h. For each tested pathogen four replicas were performed. MIC was considered the lowest concentration of CFS at which each pathogen did not grow. To determine MFC, 10 μL of the concentration corresponding to the MIC and higher concentrations were sub-cultured on PDA dishes and incubated at 25°C for 72h. Accordingly, MFC was defined as the lowest extract concentration which prevented any mycelium growth.

### 2.7 Identification of organic and phenolic acids in CFSs

To identify organic acids produced by LABs showing antifungal activity, CFSs were diluted in SDW (ratio 1/20) and injected into a high-performance liquid chromatography (HPLC) system (Agilent 1100 Series 203 HPLC System, Agilent Technologies, Palo Alto, CA, USA), equipped with a diode array and quaternary pump, using a 20 μL sample injection loop (Khosravi et al., 2015). The analytical separation was achieved with a Rezex ROA-Organic Acid H+ (8%) (150 × 7.8 mm) ion exclusion column (Phenomenex, Torrance, CA, USA) using an isocratic mobile phase of acidified water (pH 2.1) at a flow rate of 0.6 mL/min for 25 min. The chromatogram was monitored at 210 nm and results were expressed as g/L. Data were acquired by the HP209 CORE ChemStation system (Agilent Technologies, Santa Clara, CA, USA). The QuEChERS method supported the recognition of phenolic acids eliminating possible interferents before the chromatographic analysis (Brosnan et al., 2014). Extracts were prepared from CFS as reported in Dopazo et al. (2021). For the determination, a HPLC system Agilent 1200 (Agilent Technologies, Santa Clara, CA, USA) equipped with vacuum degasser, autosampler and binary pump was used. The column was a Gemini C18 (50 mm × 2 mm, 100 Å, particle size of 3-μm; Phenomenex). The mobile phases consisted of water (A) as solvent, ACN as solvent B, both acidified (0.1% formic acid), with gradient elution (0 min, 5% B; 30 min 95% B; 35 min, 5% B). Before every analysis, the column was equilibrated for 3 min. The sample (20 μL) was injected and the flow rate was 0.3 mL/min. Q-TOF-MS (6540 Agilent Ultra High Definition Accurate Mass) was used to conduct Mass spectrometry (MS) analysis equipped with an Agilent Dual Jet Stream electrospray ionisation (Dual AJS ESI) interface in negative ionisation mode following conditions: drying gas flow (N2), 8.0 L/min; nebuliser pressure, 30 psig; gas drying temperature, 350°C; capillary voltage, 3.5 kV; fragmentor voltage, 175 V; scan range, m/z 20–380. Targeted MS/MS experiments were carried out using collision energy values of 10, 20 and 40 eV. Integration and data elaboration were managed using MassHunter Qualitative Analysis software B.08.00 (Denardi-Souza et al., 2018). Results were expressed with a relative abundance (Log scale 2 to -2).

### 2.8 Analysis of the main volatile organic compounds (VOCs) from CFSs

The determination of VOCs was performed by HS-SPME (headspace solid-phase microextraction), subsequent analysis by gas chromatography coupled to mass spectrometry (GC-MS) followed the procedure of Luz et al. (2021) slightly modified.

Samples were prepared by adding 10 mL of fermented CFS in 20 mL glass vial and transferred to a 55°C bath for 45 min with constant stirring with a crystal rod throughout the whole incubation period. The SPME holder (Supelco, Bellafonte, PA, USA) contained a fused-silica fibre of 80μm x 10 mm coated with a layer of divinylbenzene–carboxen–polydimethylsiloxane (DVB/CWR/PDMS) (Agilent Technologies, Santa Clara, California, US). The fibre was introduced into the Agilent 7890A GC system coupled to an Agilent 7000A triple quadrupole mass spectrometer with an electronic impact (EI) sensor. Thermal desorption of the analytes was performed at 250°C for 10 min. Injection was performed in splitless mode. The capillary column (J&W Scientific, Folsom, CA, USA), with an HP-5MS (30 m × 0.25 mm, 0.25 μm 5% diphenyl–95% dimethylpolysiloxane) (Agilent Technologies, Santa Clara, California, USA) was used for the analysis. The carrier gas was helium (99.99%), flowing at a rate of 1 mL min^−1^. The program started at 40°C for 2 min, then increased to 160°C at 6 min; finally, the temperature was raised to 260°C at 10 °C min−1 and remained constant for 40 min. Flow in the column was transferred to an Agilent 5973 MS detector. The ion source temperature was set at 230°C, the ionizing electron energy was 70 eV and the mass range was 40–450 Da in full-scan acquisition mode. Compound identification was performed comparing their mass spectra with those recorded in an NIST Atomic Spectra Database version 1.6 (National Institute of Standards and Technologies, Gaithersburg, MD, USA), using 95% spectral similarity. Three replicas of each analysis were carried out. In addition, linear retention indices (LRI) were calculated based on the retention time of a solution of alkanes (C8-C20) tested under the same conditions as the samples and compared with the existing literature. Results were given as a percentage of each VOC in the CFS by dividing each analyte area by the total area.

### 2.9 Statistical analysis

Analysis was performed using RStudio v.1.2.5 (R). Assays were performed in triplicates, and the differences between control and treated groups were analyzed by Student’s t-test, while the differences between the groups were analyzed by one-way ANOVA test. The significance levels were set using HSD post hoc test at *p* < 0.05. VOCs were analyzed by ANOVA test and the significance levels were set at *p* ≤ 0.01. Moreover, to correlate the metabolites produced by each CFSs with the species of LAB identified, a Principal component analysis (PCA) was realised using MetaboAnalyst 5.0 software (Pang et al., 2021). The features included were log transformed and mean centred.

## 3. Results

### 3.1 Bacterial identification

Bacterial isolates (Table 1) were identified by peptide mass fingerprinting (Microflex L20) MALDI TOF MS. Identifications with a Log (score) higher than 2 were considered at species level. The isolates were identified as *Lactiplantibacillus plantarum* OV 20246 (OV8, OV9, OV16, OV17, OV20), *L. plantarum* OV 1055 (OV11), *Pediococcus pentosaceus* OV 20280 (LEC3, COR6, OT3 and OT5), *Streptococcus salivarius* OV 140417 (FR2), *Enterococcus faecium* (FR6 and COR2). Moreover, the identity of the isolates was confirmed by the full sequence of the 16S rRNA then compared to reference isolates deposited in NBCI using BlastN tool.

**Table 1:** Isolates of Lactic Acid Bacteria (LAB) sourced from oleaster (*Olea europaea oleaster*) and different cultivars of olive (*O. europaea*) in Valencia (Spain) and Calabria (Italy), respectively, and characterized in this study.

| Isolate code | Species | Host (Species and Cultivar) | Geographical Origin |
| --- | --- | --- | --- |
| OV7 | *Lactiplantibacillus plantarum* | *Olea europaea oleaster* | Villar del Arobispo, Valencia, Spain |
| OV8 | *L. plantarum* | *O. europaea oleaster* | Villar del Arobispo, Valencia, Spain |
| OV9 | *L. plantarum* | *O. europaea oleaster* | Villar del Arobispo, Valencia, Spain |
| OV11 | *L. plantarum* | *O. europaea oleaster* | Higueruelas, Valencia, Spain |
| OV16 | *L. plantarum* | *O. europaea oleaster* | Higueruelas, Valencia, Spain |
| OV17 | *L. plantarum* | *O. europaea oleaster* | Higueruelas, Valencia, Spain |
| OV20 | *L. plantarum* | *O. europaea oleaster* | Higueruelas, Valencia, Spain |
| FR2 | *Streptococcus salivarius* | *O. europaea* 'Frantoio' | Mirto-Crosia, Cosenza, Italy |
| FR6 | *Enterococcus faecium* | *O. europaea* 'Frantoio' | Mirto-Crosia, Cosenza, Italy |
| LEC3 | *Pediococcus pentosaceus* | *O. europaea* 'Leccino' | Mirto-Crosia, Cosenza, Italy |
| OT3 | *P. pentosaceus* | *O. europaea* 'Ottobratica' | Mirto-Crosia, Cosenza, Italy |
| OT5 | *P. pentosaceus* | *O. europaea* 'Ottobratica' | Mirto-Crosia, Cosenza, Italy |
| COR2 | *E. faecium* | *O. europaea* 'Coratina' | Mirto-Crosia, Cosenza, Italy |
| COR3 | *P. pentosaceus* | *O. europaea* 'Coratina' | Mirto-Crosia, Cosenza, Italy |
| COR5 | *P. pentosaceus* | *O. europaea* 'Coratina' | Mirto-Crosia, Cosenza, Italy |
| COR6 | *P. pentosaceus* | *O. europaea* 'Coratina' | Mirto-Crosia, Cosenza, Italy |

### 3.2 Qualitative assay of antifungal activity in solid medium

In *in vitro* assays, all 16 LABs showed antifungal activity. However, there was a consistent variability in the inhibitory efficacy among bacterial isolates against the diverse fungi *s*.*l*. tested. In the overlay test (Table 2), both isolates of *E. faecium* (FR6 and COR2) showed no antifungal activity and none of the bacterial isolates inhibited the mycelium growth of *A. flavus* and *Plenodomus tracheiphilus*. The most noticeable inhibitory activity was shown by isolates of *L. plantarum* and *P. pentosaceus*. Isolates of these two bacterial species exerted a relevant inhibitory activity versus *Fusarium oxysporum* and species of *Colletotrichum* with inhibition halos in most cases larger than 10 mm. Large (diameter > 10 mm) and medium (diameter 8÷10 mm) inhibition halos were also observed in tests against *P. nordicum*. The inhibitory activity versus other *Penicillium* species was less consistent. All *L. plantarum, P. pentosaceus* and *S. salivarius* isolates showed inhibitory activity against *A. alternata*. However, the most effective were isolates OV8, OV17, COR3 and LEC3, whose inhibition halos were medium (diameter 8-10 mm) to large (diameter > 10 mm) in size. In tests against *Phytophthora* isolates the maximum inhibitory activity (mean diameter of inhibition halo > 10 mm) was shown by the bacterial isolate COR6 (*P. pentosaceus*). The inhibition halos induced by the other *L. plantarum, P. pentosaceus* and *S. salivarius* isolates were medium (diameter 8-10 mm) to low (diameter < 8mm) in size.

**Table 2.** Antifungal activity of lactic acid bacteria (LABs) against *Colletotrichum, Penicillium, Aspergillus, Fusarium, Alternaria, Plenodomus* and *Phytophthora* species in the culture overlay test. The antifungal activity was represented as follows: (+) means of inhibition zone between the well and fungal growth 8 mm, (++) means of inhibition zone between the well and fungal growth 8–10 mm, (+++) means of inhibition zone between the well and fungal growth > 10 mm. The well radius was 5 mm.

| Pathogen | Isolate code | Lactic Acid Bacteria (LABs) |  |  |  |  |  |  |  |  |  |  |  |  |  |  |  |
| --- | --- | --- | --- | --- | --- | --- | --- | --- | --- | --- | --- | --- | --- | --- | --- | --- | --- |
|  |  | OV7 | OV8 | OV9 | OV11 | OV16 | OV17 | OV20 | COR2 | COR3 | COR5 | COR6 | OT3 | OT5 | LEC3 | FR2 | FR6 |
| *Alternaria alternata* | ITEM 8121 | + | +++ | + | + | + | ++ | + | - | ++ | + | + | + | + | ++ | + | - |
| *A. alternata* | CECT 646 | + | +++ | + | + | + | +++ | + | - | ++ | + | + | + | + | +++ | + | - |
| *Aspergillus flavus* | ISPA<br>8111 | - | - | - | - | - | - | - | - | - | - | - | - | - | - | - | - |
| *Colletotrichum*<br>*acutatum* | C9D2C | +++ | +++ | +++ | +++ | +++ | ++ | +++ | - | ++ | + | +++ | ++ | +++ | +++ | + | - |
| *C. acutatum* | UWS<br>149 | +++ | +++ | +++ | +++ | +++ | + | +++ | - | ++ | + | +++ | ++ | +++ | +++ | + | - |
| *C. fiorinae* | C15D6A | ++ | +++ | +++ | ++ | +++ | + | + | - | + | ++ | +++ | ++ | +++ | +++ | ++ | - |
| *C. fiorinae* | ER2147 | ++ | ++ | +++ | ++ | +++ | + | + | - | + | ++ | ++ | ++ | +++ | ++ | ++ | - |
| *C.*<br>*gloeosporioides* | RD9B | ++ | +++ | + | + | +++ | +++ | + | - | + | - | ++ | + | ++ | ++ | ++ | - |
| *C.*<br>*gloeosporioides* | C2 | ++ | ++ | + | + | +++ | ++ | + | - | ++ | - | ++ | + | ++ | ++ | ++ | - |
| *C. nymphaeae* | RB012 | ++ | +++ | ++ | ++ | ++ | +++ | - | - | + | +++ | +++ | ++ | +++ | ++ | ++ | - |
| *C. nymphaeae* | RB428 | ++ | +++ | ++ | ++ | ++ | +++ | - | - | + | +++ | +++ | ++ | +++ | ++ | ++ | - |
| *Fusarium*<br>*oxysporum* | COAL<br>68 | +++ | +++ | ++ | +++ | +++ | +++ | ++ | - | + | + | ++ | + | ++ | +++ | ++ | - |
| *Penicillium*<br>*digitatum* | P1PP0 | + | + | ++ | - | + | + | + | - | + | ++ | - | + | + | - | + | - |
| *P. digitatum* | N2F1 | + | + | + | - | + | + | + | - | + | ++ | - | + | + | - | + | - |
| *P. expansum* | COAL<br>95 | + | + | + | + | + | ++ | ++ | - | - | - | - | - | + | - | - | + |
| *P. nordicum* | CECT<br>2320 | ++ | ++ | +++ | + | +++ | - | +++ | - | +++ | + | +++ | + | + | + | + | - |
| *Phytophthora*<br>*nicotianae* | P4K3F8 | ++ | ++ | ++ | ++ | + | + | ++ | - | + | ++ | ++ | + | + | ++ | + | - |
| *Ph. nicotianae* | T-2K1A | ++ | ++ | ++ | ++ | + | + | ++ | - | + | ++ | +++ | + | + | ++ | ++ | - |
| *Ph. oleae* | V-2K10A | ++ | ++ | ++ | + | + | + | ++ | - | + | + | ++ | + | + | ++ | + | - |
| *Plenodomus*<br>*tracheiphilus* | Pt.2 | - | - | - | - | - | - | - | - | - | - | - | - | - | - | - | - |

In the agar diffusion test (Table 3) all LABs, including the isolates of *E. faecium* (FR6 and COR2), showed antifungal activity. With few exceptions, results of this test agreed with those of the culture overlay test. None of the CFSs, in fact, inhibited the mycelial growth of *P. tracheiphilus* and CFSs of most bacterial isolates showed no antifungal activity against *A. flavus*. Only CFSs of bacterial isolates OV8, OV20 and FR6 inhibited the growth of this fungus and in any case the mean diameter of inhibition halos was < 8 mm. It was confirmed a noticeable inhibitory activity of *L. plantarum* and *P. pentosaceus* on *Colletotrichum* species and *F. oxysporum*. Also, the fermentate of FR6 isolate (*E. faecium*) showed a high inhibitory activity against *Colletotrichum* species and *F. oxysporum*. Although this LAB did not show any antifungal activity in the culture overlay test, in the agar diffusion tests its fermentate exerted a notable inhibition versus *Colletotrichum* species and *F. oxysporum* (diameter of inhibition halos >10 mm). Consistently with the results of the culture overlay test, *P. nordicum* was remarkably inhibited by LAB fermentates. Lab isolates with the highest inhibitory effect (diameter of inhibition halos >10 mm) were OV7, OV8 and OV11, all identified as *L. plantarum*. Inhibition halos of all other *Penicillium* species never exceeded 10 mm in diameter, in most cases they were <8 mm in diameter or no inhibitory effect was detected. CFSs of all bacterial species showed an inhibitory effect on one or both isolates of *A. alternata*, with the only exception of the CFS of COR 6 *P. pentosaceus* isolate. However, the diameter of inhibition halos was always <10 mm. CFSs of three *L. plantarum* isolates (OV8, OV11 and OV20) showed a relevant inhibitory activity against *P. nicotianae* and *P. olea*e (diameter of inhibition halos 8-10 mm), while CFSs of other bacterial isolates exerted only a low inhibitory activity (diameter of inhibition halos <8mm) or did not exert any inhibitory activity on these oomycetes.

**Table 3.** Antifungal activity of sixteen cell-free supernatants (CFS) of lactic acid bacteria (LABs) against *Colletotrichum, Penicillium, Aspergillus, Fusarium, Alternaria, Plenodomus* and *Phytophthora* species in the diffusion agar test. The antifungal activity was represented as follows: (+) means 8 mm of inhibition zone between the well and fungal growth, (++) means 8–10 mm of inhibition zone between the well and fungal growth, (+++) means > 10 mm of inhibition zone between the well and fungal growth. The well radius was 5 mm. A CFS concentration of 400 mg/m was used. MRS medium was used as control.

| Pathogen | Isolate code | CFS of Lactic Acid Bacteria (LABs) |  |  |  |  |  |  |  |  |  |  |  |  |  |  |  |
| --- | --- | --- | --- | --- | --- | --- | --- | --- | --- | --- | --- | --- | --- | --- | --- | --- | --- |
|  |  | OV7 | OV8 | OV9 | OV11 | OV16 | OV17 | OV20 | COR2 | COR3 | COR5 | COR6 | OT3 | OT5 | LEC3 | FR2 | FR6 |
| *Alternaria alternata* | CECT 646 | ++ | ++ | + | + | - | + | ++ | ++ | - | + | - | + | - | + | ++ | + |
| *A. alternata* | ITEM 8121 | ++ | ++ | + | ++ | + | + | ++ | ++ | + | + | - | - | + | + | + | + |
| *Aspergillus flavus* | ITEM 8111 | - | + | - | - | - | - | + | - | - | - | - | - | - | - | - | + |
| *Colletotrichum acutatum* | UWS149 | +++ | +++ | +++ | ++ | ++ | +++ | +++ | ++ | +++ | ++ | ++ | ++ | ++ | ++ | + | ++ |
| *C. acutatum* | C9D2C | +++ | +++ | +++ | ++ | ++ | +++ | +++ | ++ | +++ | ++ | ++ | ++ | ++ | ++ | + | ++ |
| *C. fiorinae* | ER2147 | ++ | +++ | +++ | ++ | ++ | +++ | +++ | ++ | + | ++ | + | ++ | + | - | ++ | +++ |
| *C. fiorinae* | C15D6A | +++ | +++ | +++ | +++ | ++ | +++ | +++ | ++ | + | ++ | + | ++ | + | + | ++ | +++ |
| *C. gloeosporioides* | C2 | + | ++ | ++ | +++ | ++ | + | +++ | ++ | + | + | + | + | ++ | + | + | ++ |
| *C. gloeosporioides* | RD9B | ++ | +++ | ++ | +++ | ++ | + | +++ | ++ | + | ++ | + | ++ | ++ | + | + | ++ |
| *C. godetiae* | OLP10 | ++ | ++ | + | +++ | ++ | + | +++ | ++ | + | + | + | ++ | +++ | + | ++ | ++ |
| *C. godetiae* | OLP12 | ++ | ++ | + | +++ | ++ | + | +++ | ++ | + | + | + | ++ | +++ | + | ++ | ++ |
| *C. nymphaeae* | RB428 | ++ | +++ | ++ | ++ | ++ | ++ | +++ | ++ | + | ++ | + | ++ | ++ | ++ | ++ | +++ |
| *C. nymphaeae* | RB012 | ++ | +++ | ++ | ++ | ++ | ++ | +++ | ++ | + | ++ | + | ++ | ++ | ++ | ++ | +++ |
| *Fusarium oxysporum* | COAL 68 | +++ | +++ | ++ | ++ | ++ | ++ | +++ | + | ++ | + | - | ++ | ++ | + | ++ | +++ |
| *Penicillium digitatum* | N2F1 | + | + | + | + | + | + | + | ++ | - | + | + | - | - | - | - | - |
| *P. digitatum* | P1PP0 | + | + | + | + | + | + | + | + | - | + | + | - | - | - | - | - |
| *P. expansum* | COAL 95 | + | + | + | + | + | ++ | ++ | - | - | - | - | - | + | - | - | + |
| *P. nordicum* | CECT 2320 | +++ | +++ | ++ | +++ | ++ | ++ | ++ | ++ | ++ | + | + | + | ++ | + | ++ | ++ |
| *Phytophthora nicotianae* | T-2K1A | + | ++ | + | ++ | + | - | ++ | + | - | - | - | + | - | - | - | + |
| *Ph. nicotianae* | P4K3F8 | + | ++ | + | ++ | + | - | ++ | + | - | - | - | - | - | - | - | - |
| *Ph. oleae* | V-2K10A | + | ++ | + | ++ | + | - | ++ | - | - | - | - | - | - | - | - | - |
| *Plenodomus tracheiphilus* | Pt 2 | - | - | - | - | - | - | - | - | - | - | - | - | - | - | - | - |

### 3.3 Quantitative assay of antifungal activity in liquid medium

The results of assays, aimed at determining the values of MIC and MFC and thus quantifying the antifungal activity of CFSs of LABs, are summarized in Table 4. In general, the values of MIC and MFC were in the same order of magnitude and in most cases overlapped. Consistently with the results of both the culture overlay and the diffusion tests, the CFSs that showed high antifungal activity, i.e. low values of MIC and MFC, included the CFSs of five LAB isolates, OV7, OV8, OV9, OV11 and OV20, identified as *L. plantarum*), LEC3 (*P. pentosaceus*) and FR6 (*E. faecium*). Particularly, the MIC and MFC values of OV7 bacterial isolate were in the range of 12.5 to 100 g/L. The highest MIC and MFC values (100 g/L) for CFS of this *L. plantarum* isolate were recorded against *Ph. nicotianae*. Conversely, the lowest value (12.5 g/L) was recorded in assays versus isolates of *Colletotrichum*. However, differently from the other isolates of *Colletotrichum*, both MIC and MFC values against C2 and RD9B isolates of *C. gloeosporioides*, recovered from distinct host plants, were relatively high, i.e., 100 g/L (for both MIC and MFC against C2 isolate) and 50 g/L (for both MIC and MFC against isolate RD9B). Isolate OV8 of *L. plantarum*, which in general showed a comparable antifungal activity to that of isolate OV7, was more effective in inhibiting the mycelium growth of both C2 and RD9B isolates of *C. gloeosporioides*. MIC and MFC values against these two fungus isolates were 25 versus 12.5 g/L and 50 versus 50, respectively. Moreover, while MIC and MFC values against *Phytophthora* isolates ranged between 25 and 50 g/L for the LAB isolate OV8, they ranged between 10 and 100 g/L for isolate OV7. The CFS of *E. faecium* FR6 isolate, which showed no antifungal activity in the overlay test, but showed a consistent inhibitory activity in the agar diffusion test, inhibited all tested pathogens at concentrations ranging from 12.5 to 50 g/L for MIC and 25 to 50 g/L for MFC. MIC and MFC values for the CFS of this LAB isolate against isolates of *Phytophthora* were relatively high (100 g/L). Finally, FR6 isolate showed a noticeable antifungal activity against *P. nordicum* (MIC and MFC 12.5 and 25 g/L, respectively) and a relatively low antifungal activity against *P. expansum* and *P. digitatum* (MIC and MFC values between 50 and 100 g/L). The CFS of *P. pentosaceous* isolate LEC 3 showed a consistent inhibitory activity against all fungi and oomycetes tested, with MIC and MFC values ranging from 25 and 50 g/L.

**Table 4.**
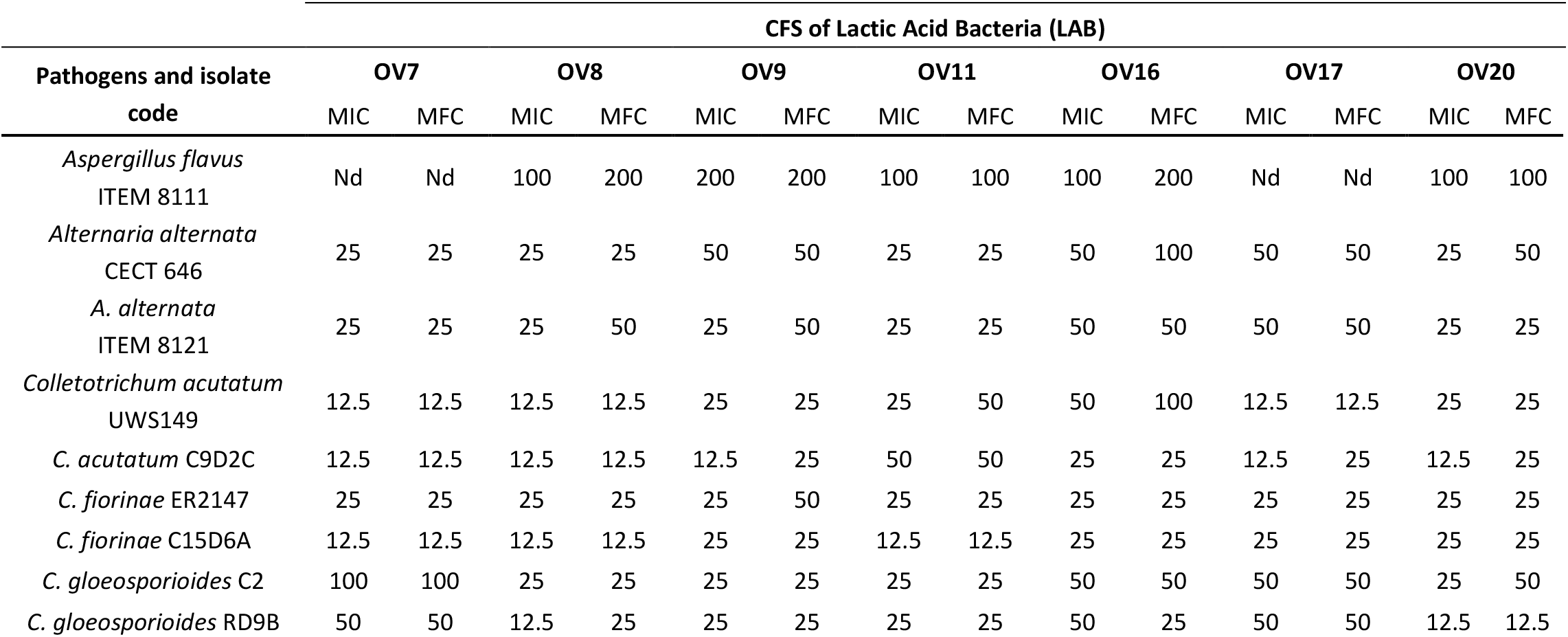
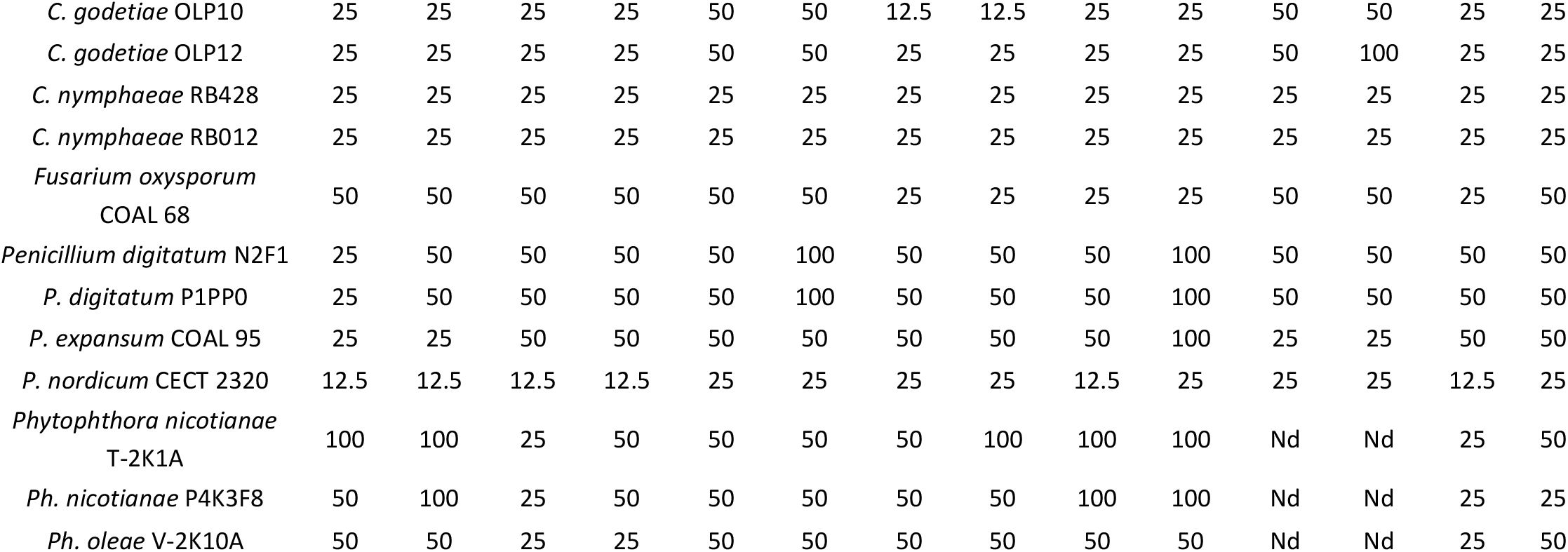
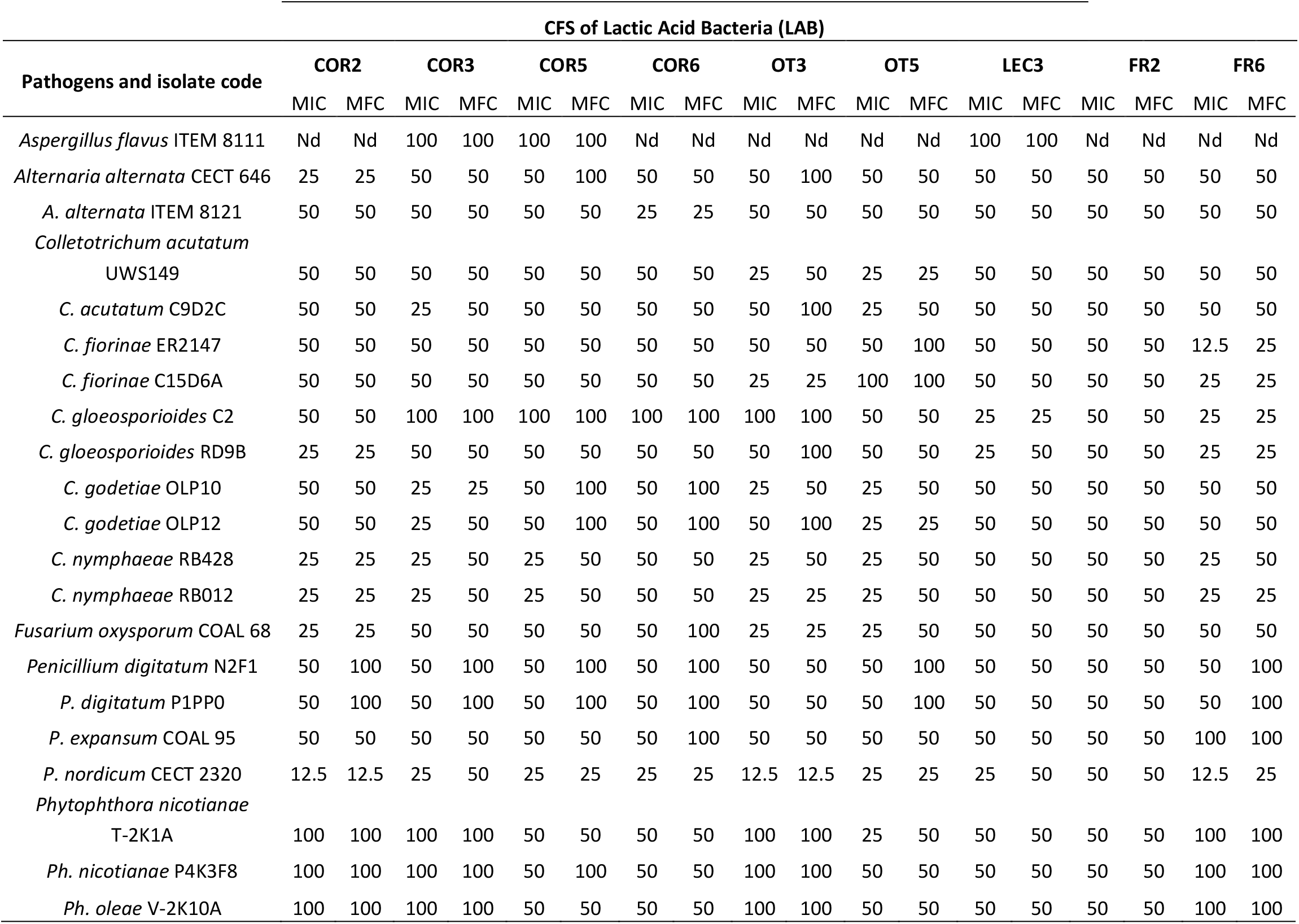
Minimum Inhibitory Concentration (MIC) and Minimum Fungicidal Concentration (MFC) of lyophilised CFSs of lactic acid bacteria expressed in g/L. nd = not detected.

| Pathogens and isolate<br>code | CFS of Lactic Acid Bacteria (LAB) |  |  |  |  |  |  |  |  |  |  |  |  |  |
| --- | --- | --- | --- | --- | --- | --- | --- | --- | --- | --- | --- | --- | --- | --- |
|  | OV7 |  | OV8 |  | OV9 |  | OV11 |  | OV16 |  | OV17 |  | OV20 |  |
|  | MIC | MFC | MIC | MFC | MIC | MFC | MIC | MFC | MIC | MFC | MIC | MFC | MIC | MFC |
| *Aspergillus flavus*<br>ITEM 8111 | Nd | Nd | 100 | 200 | 200 | 200 | 100 | 100 | 100 | 200 | Nd | Nd | 100 | 100 |
| *Alternaria alternata*<br>CECT 646 | 25 | 25 | 25 | 25 | 50 | 50 | 25 | 25 | 50 | 100 | 50 | 50 | 25 | 50 |
| *A. alternata*<br>ITEM 8121 | 25 | 25 | 25 | 50 | 25 | 50 | 25 | 25 | 50 | 50 | 50 | 50 | 25 | 25 |
| *Colletotrichum acutatum*<br>UWS149 | 12.5 | 12.5 | 12.5 | 12.5 | 25 | 25 | 25 | 50 | 50 | 100 | 12.5 | 12.5 | 25 | 25 |
| *C. acutatum* C9D2C | 12.5 | 12.5 | 12.5 | 12.5 | 12.5 | 25 | 50 | 50 | 25 | 25 | 12.5 | 25 | 12.5 | 25 |
| *C. fiorinae* ER2147 | 25 | 25 | 25 | 25 | 25 | 50 | 25 | 25 | 25 | 25 | 25 | 25 | 25 | 25 |
| *C. fiorinae* C15D6A | 12.5 | 12.5 | 12.5 | 12.5 | 25 | 25 | 12.5 | 12.5 | 25 | 25 | 25 | 25 | 25 | 25 |
| *C. gloeosporioides* C2 | 100 | 100 | 25 | 25 | 25 | 25 | 25 | 25 | 50 | 50 | 50 | 50 | 25 | 50 |
| *C. gloeosporioides* RD9B | 50 | 50 | 12.5 | 25 | 25 | 25 | 25 | 25 | 50 | 25 | 50 | 50 | 12.5 | 12.5 |
| *C. godetiae* OLP10 | 25 | 25 | 25 | 25 | 50 | 50 | 12.5 | 12.5 | 25 | 25 | 50 | 50 | 25 | 25 |
| *C. godetiae* OLP12 | 25 | 25 | 25 | 25 | 50 | 50 | 25 | 25 | 25 | 25 | 50 | 100 | 25 | 25 |
| *C. nymphaeae* RB428 | 25 | 25 | 25 | 25 | 25 | 25 | 25 | 25 | 25 | 25 | 25 | 25 | 25 | 25 |
| *C. nymphaeae* RB012 | 25 | 25 | 25 | 25 | 25 | 25 | 25 | 25 | 25 | 25 | 25 | 25 | 25 | 25 |
| *Fusarium oxysporum*<br>COAL 68 | 50 | 50 | 50 | 50 | 50 | 50 | 25 | 25 | 25 | 25 | 50 | 50 | 25 | 50 |
| *Penicillium digitatum* N2F1 | 25 | 50 | 50 | 50 | 50 | 100 | 50 | 50 | 50 | 100 | 50 | 50 | 50 | 50 |
| *P. digitatum* P1PP0 | 25 | 50 | 50 | 50 | 50 | 100 | 50 | 50 | 50 | 100 | 50 | 50 | 50 | 50 |
| *P. expansum* COAL 95 | 25 | 25 | 50 | 50 | 50 | 50 | 50 | 50 | 50 | 100 | 25 | 25 | 50 | 50 |
| *P. nordicum* CECT 2320 | 12.5 | 12.5 | 12.5 | 12.5 | 25 | 25 | 25 | 25 | 12.5 | 25 | 25 | 25 | 12.5 | 25 |
| *Phytophthora nicotianae*<br>T-2K1A | 100 | 100 | 25 | 50 | 50 | 50 | 50 | 100 | 100 | 100 | Nd | Nd | 25 | 50 |
| *Ph. nicotianae* P4K3F8 | 50 | 100 | 25 | 50 | 50 | 50 | 50 | 50 | 100 | 100 | Nd | Nd | 25 | 25 |
| *Ph. oleae* V-2K10A | 50 | 50 | 25 | 25 | 50 | 50 | 50 | 50 | 50 | 50 | Nd | Nd | 25 | 50 |

**Table 4.** (Continue)
| Pathogens and isolate code | CFS of Lactic Acid Bacteria (LAB) |  |  |  |  |  |  |  |  |  |  |  |  |  |  |  |  |  |
| --- | --- | --- | --- | --- | --- | --- | --- | --- | --- | --- | --- | --- | --- | --- | --- | --- | --- | --- |
|  | COR2 |  | COR3 |  | COR5 |  | COR6 |  | OT3 |  | OT5 |  | LEC3 |  | FR2 |  | FR6 |  |
|  | MIC | MFC | MIC | MFC | MIC | MFC | MIC | MFC | MIC | MFC | MIC | MFC | MIC | MFC | MIC | MFC | MIC | MFC |
| *Aspergillus flavus* ITEM 8111 | Nd | Nd | 100 | 100 | 100 | 100 | Nd | Nd | Nd | Nd | Nd | Nd | 100 | 100 | Nd | Nd | Nd | Nd |
| *Alternaria alternata* CECT 646 | 25 | 25 | 50 | 50 | 50 | 100 | 50 | 50 | 50 | 100 | 50 | 50 | 50 | 50 | 50 | 50 | 50 | 50 |
| *A. alternata* ITEM 8121 | 50 | 50 | 50 | 50 | 50 | 50 | 25 | 25 | 50 | 50 | 50 | 50 | 50 | 50 | 50 | 50 | 50 | 50 |
| *Colletotrichum acutatum*<br>UWS149 | 50 | 50 | 50 | 50 | 50 | 50 | 50 | 50 | 25 | 50 | 25 | 25 | 50 | 50 | 50 | 50 | 50 | 50 |
| *C. acutatum* C9D2C | 50 | 50 | 25 | 50 | 50 | 50 | 50 | 50 | 50 | 100 | 25 | 50 | 50 | 50 | 50 | 50 | 50 | 50 |
| *C. fiorinae* ER2147 | 50 | 50 | 50 | 50 | 50 | 50 | 50 | 50 | 50 | 50 | 50 | 100 | 50 | 50 | 50 | 50 | 12.5 | 25 |
| *C. fiorinae* C15D6A | 50 | 50 | 50 | 50 | 50 | 50 | 50 | 50 | 25 | 25 | 100 | 100 | 50 | 50 | 50 | 50 | 25 | 25 |
| *C. gloeosporioides* C2 | 50 | 50 | 100 | 100 | 100 | 100 | 100 | 100 | 100 | 100 | 50 | 50 | 25 | 25 | 50 | 50 | 25 | 25 |
| *C. gloeosporioides* RD9B | 25 | 25 | 50 | 50 | 50 | 50 | 50 | 50 | 50 | 100 | 50 | 50 | 25 | 50 | 50 | 50 | 25 | 25 |
| *C. godetiae* OLP10 | 50 | 50 | 25 | 25 | 50 | 100 | 50 | 100 | 25 | 50 | 25 | 50 | 50 | 50 | 50 | 50 | 50 | 50 |
| *C. godetiae* OLP12 | 50 | 50 | 25 | 50 | 50 | 100 | 50 | 100 | 50 | 100 | 25 | 25 | 50 | 50 | 50 | 50 | 50 | 50 |
| *C. nymphaeae* RB428 | 25 | 25 | 25 | 50 | 25 | 50 | 50 | 50 | 25 | 50 | 25 | 50 | 50 | 50 | 50 | 50 | 25 | 50 |
| *C. nymphaeae* RB012 | 25 | 25 | 25 | 50 | 25 | 50 | 50 | 50 | 25 | 25 | 25 | 50 | 50 | 50 | 50 | 50 | 25 | 25 |
| *Fusarium oxysporum* COAL 68 | 25 | 25 | 50 | 50 | 50 | 50 | 50 | 100 | 25 | 25 | 25 | 50 | 50 | 50 | 50 | 50 | 50 | 50 |
| *Penicillium digitatum* N2F1 | 50 | 100 | 50 | 100 | 50 | 100 | 50 | 100 | 50 | 50 | 50 | 100 | 50 | 50 | 50 | 50 | 50 | 100 |
| *P. digitatum* P1PP0 | 50 | 100 | 50 | 100 | 50 | 100 | 50 | 100 | 50 | 50 | 50 | 100 | 50 | 50 | 50 | 50 | 50 | 100 |
| *P. expansum* COAL 95 | 50 | 50 | 50 | 50 | 50 | 50 | 50 | 100 | 50 | 50 | 50 | 50 | 50 | 50 | 50 | 50 | 100 | 100 |
| *P. nordicum* CECT 2320 | 12.5 | 12.5 | 25 | 50 | 25 | 25 | 25 | 25 | 12.5 | 12.5 | 25 | 25 | 25 | 50 | 50 | 50 | 12.5 | 25 |
| *Phytophthora nicotianae*<br>T-2K1A | 100 | 100 | 100 | 100 | 50 | 50 | 50 | 50 | 100 | 100 | 25 | 50 | 50 | 50 | 50 | 50 | 100 | 100 |
| *Ph. nicotianae* P4K3F8 | 100 | 100 | 100 | 100 | 50 | 100 | 50 | 50 | 100 | 100 | 50 | 50 | 50 | 50 | 50 | 50 | 100 | 100 |
| *Ph. oleae* V-2K10A | 100 | 100 | 100 | 100 | 50 | 50 | 50 | 50 | 100 | 100 | 50 | 50 | 50 | 50 | 50 | 50 | 100 | 100 |

General trends, confirming in part the results of culture overlay and agar diffusion tests, were observed. The antifungal activity against *A. flavus* of CFSs of all LABs was very low (values of both MIC and MFC ≥ 100 g/L) or in many cases out of the detectable range. CFSs of all LABs showed a clear antifungal activity against *A. alternata* and *F. oxysporum*, with MIC values ranging between 25 and 50 g/L. The antifungal activity of CFSs against *Colletotrichum* species was even more pronounced. With few exceptions, including the two above mentioned isolates of *C. gloeosporioides* (C2 and RD9B) and isolates of *C. godetiae*, MIC values against isolates of this fungal genus ranged between 12.5 and 50 g/L. MIC values of CFSs against *P. nordicum* were in the same range, confirming the high sensitivity of this isolate to the inhibitory effects of LABs. However, the antifungal activity of CFSs against the other two *Penicillium* species, *P. digitatum* and *P. expansum*, was less remarkable and lower compared to species of *Colletotrichum*. MIC values for the last two species of *Penicillium* ranged between 25 and 50 g/L. In general, isolates of *Phytophthora* showed a lower sensitivity to all tested LAB fermentates compared to species of *Colletotrichum* and *Penicillium*. MIC and MFC values for these oomycetes were in the range between 25 and 100 g/L and in a few cases exceeded 100 g/l.

### 3.4 Ability of selected bacteria to produce lactic acid, phenolic acids and VOC

Overall, an organic acid (lactic acid) and 14 phenolic acids were characterized and quantified in CFSs of LABs isolates. Comparing the data by ANOVA there were significant differences in metabolite profile among the 16 bacterial isolates. All isolates produced lactic acid, as expected, with values ranging from 18.4 to 32.8 g/L (Table 5). The highest concentrations of lactic acid were recorded in CFSs of OV8, OV7, OV9, OV11, OV17 and OV20 *L. plantarum* isolates. Conversely, the lowest concentration of lactic acid (18.4 ± 1.6 g/L) was found in CFS of FR2 isolate (*S. salivarius*) (Table 5).

**Table 5.** Quantification of lactic acid (g/L) produced in CFSs by bacterial strains isolated from olives. Data are means ± standard deviation. Letters indicate statistically significant differences using one-way ANOVA Tukey HSD post hoc test (*p*< 0.05).

| Isolate code | Strain | Lactic acid (g/L) |
| --- | --- | --- |
| OV7 | *Lactiplantibacillus plantarum* | $31.4 \pm 1.0^{ef}$ |
| OV8 | *L. plantarum* | $32.8 \pm 0.8^f$ |
| OV9 | *L. plantarum* | $32.6 \pm 1.3^f$ |
| OV11 | *L. plantarum* | $31.6 \pm 0.3^{ef}$ |
| OV16 | *L. plantarum* | $29.7 \pm 1.1^{def}$ |
| OV17 | *L. plantarum* | $31.6 \pm 0.7^{ef}$ |
| OV20 | *L. plantarum* | $30.6 \pm 0.4^{ef}$ |
| LEC3 | *Pediococcus pentosaceus* | $25.9 \pm 0.3^{bcd}$ |
| OT3 | *P. pentosaceus* | $27.6 \pm 1.3^{bcde}$ |
| OT5 | *P. pentosaceus* | $23.7 \pm 1.4^b$ |
| COR3 | *P. pentosaceus* | $28 \pm 2.4^{cde}$ |
| COR5 | *P. pentosaceus* | $26 \pm 2.7^{bcd}$ |
| COR6 | *P. pentosaceus* | $24 \pm 0.6^{bc}$ |
| FR2 | *Streptococcus salivarius* | $18.4 \pm 1.6^a$ |
| FR6 | *Enterococcus faecium* | $23.4 \pm 1.6^b$ |
| COR2 | *E. faecium* | $26 \pm 2.4^{cde}$ |

The phenolic acids detected and quantified in the CFSs were 1-2-dihydroxybenzene, 3-(4-hydroxy-3- methoxyphenyl) propionic, 3-4-dihydroxyhydrocinnamic, benzoic acid, caffeic acid, DL-3-phenyllactic acid (PLA), ferulic acid, hydroxicinnamic acid, P- coumaric acid, salicilic acid, sinapic acid, syringic acid, vanillic acid and vanillin. Only *L. plantarum* OV8 strain produced high values of all acids (Figure 1).

**Figure 1.**
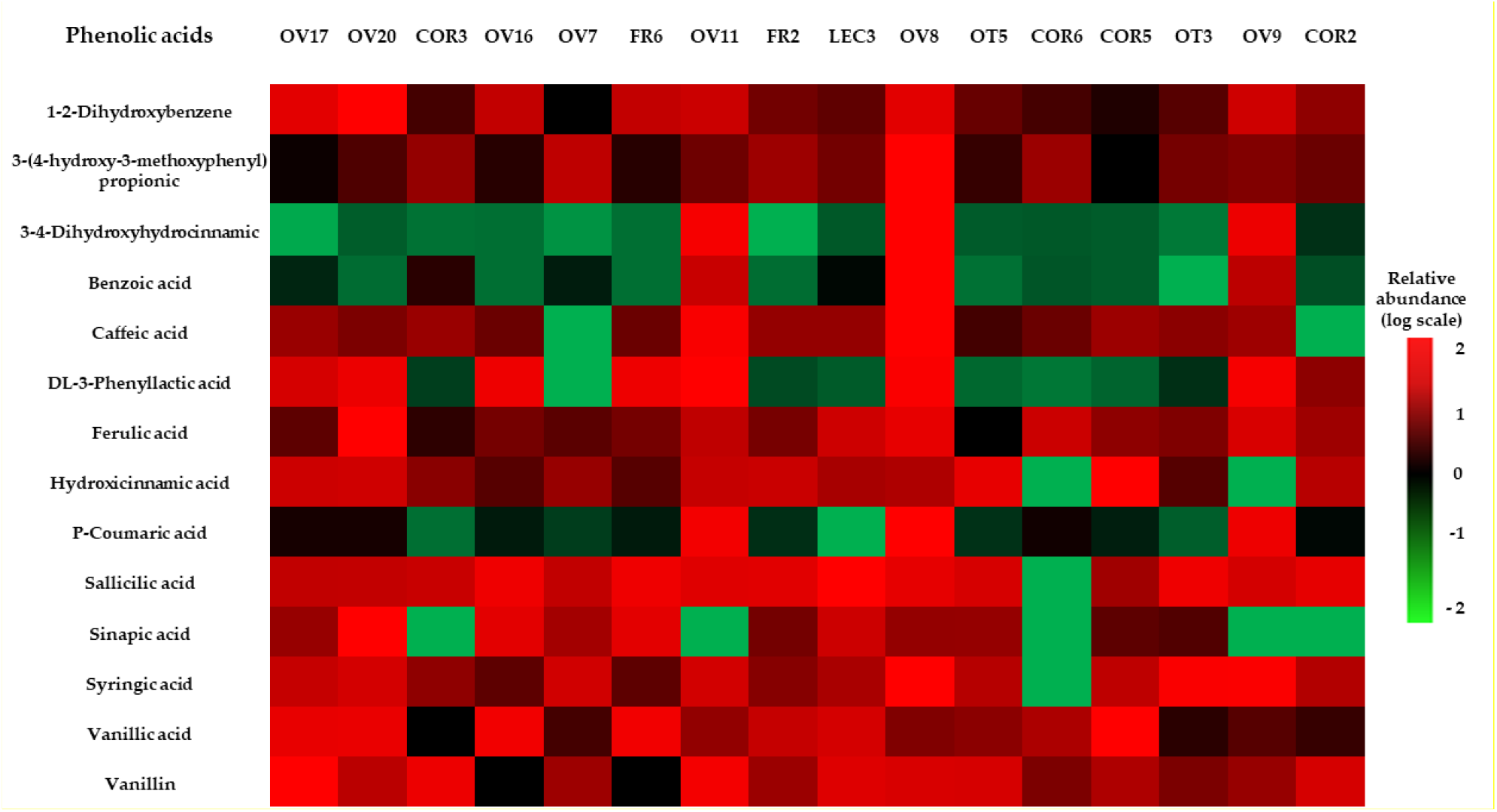
Heat map of phenolic acid produced by tested bacterial strains. Colors are based on relative abundance (logarithmic scale) of produced metabolites, independently for each bacterium: red represents high abundance and green low abundance.

PLA was recorded with a high relative abundance (positive log scale) in CFS s of OV8, OV9, OV11, OV16, OV20 (*L. plantarum*) and FR6 (*E. faecium*). Moreover, high relative abundance of benzoic and 3-4- dydroxyhydrocinnamic acids were found in CFSs of *L. plantarum* OV11, OV8 and OV9 isolates. Benzoic acid was not recorded in CFSs of OV7, OV17, COR3 and LEC3 *L. plantarum* and *P. pentosaceus* isolates. Caffeic acid, vanillic acid and vanillin were produced in abundance by all LABS, except for OV 16 and FR 6 for vanillin, COR 3 for vanillic acid and OV17 for caffeic acid.

The volatile metabolites extracted from CFSs were characterized using the HS-SPME/GC-MS technique. A total of 26 VOCs from CFSs and the control (MRS-B) were detected and quantified. The classes identification of all detected VOCs is reported in Table S1. The VOCs were divided into seven classes based on their chemical properties: acids (0% - 15.7%), alcohols (0% - 13.6%), aldehydes (0% - 3.2 %), alkanes (4.1% - 15.2%), alkenes (24.7% - 40.3%), ketones (6.1%- 42.4%) and esters (0 – 5.4%) (Table 6).

**Table 6.**
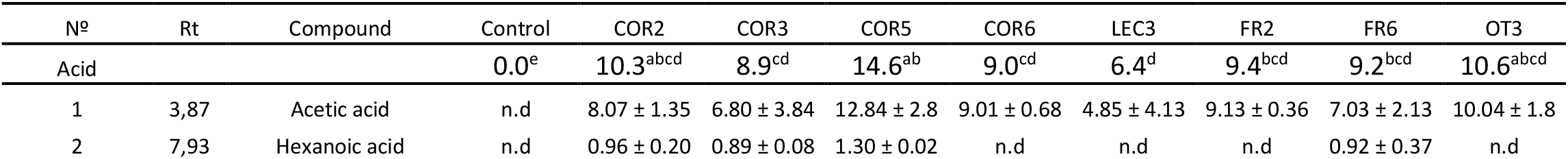
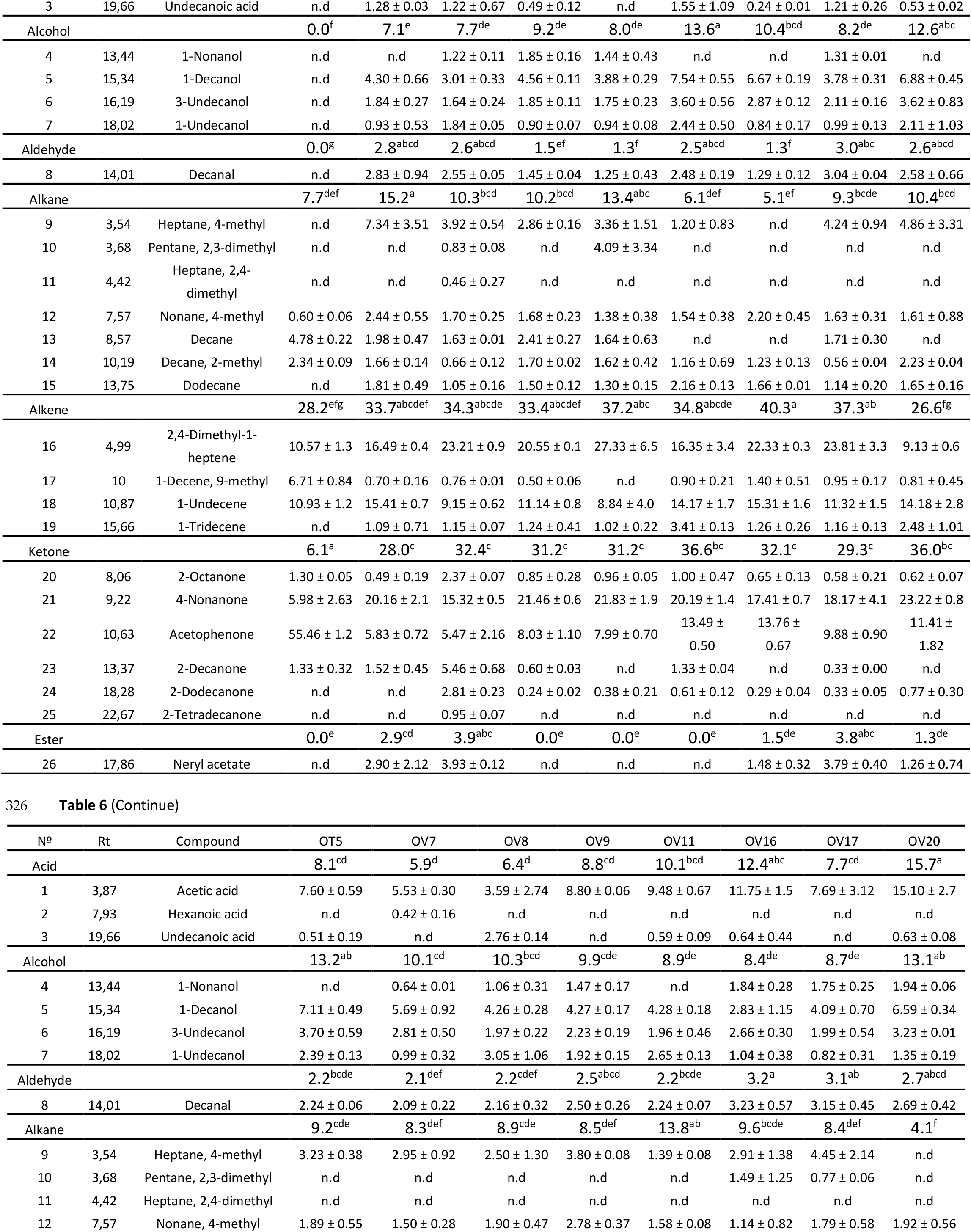
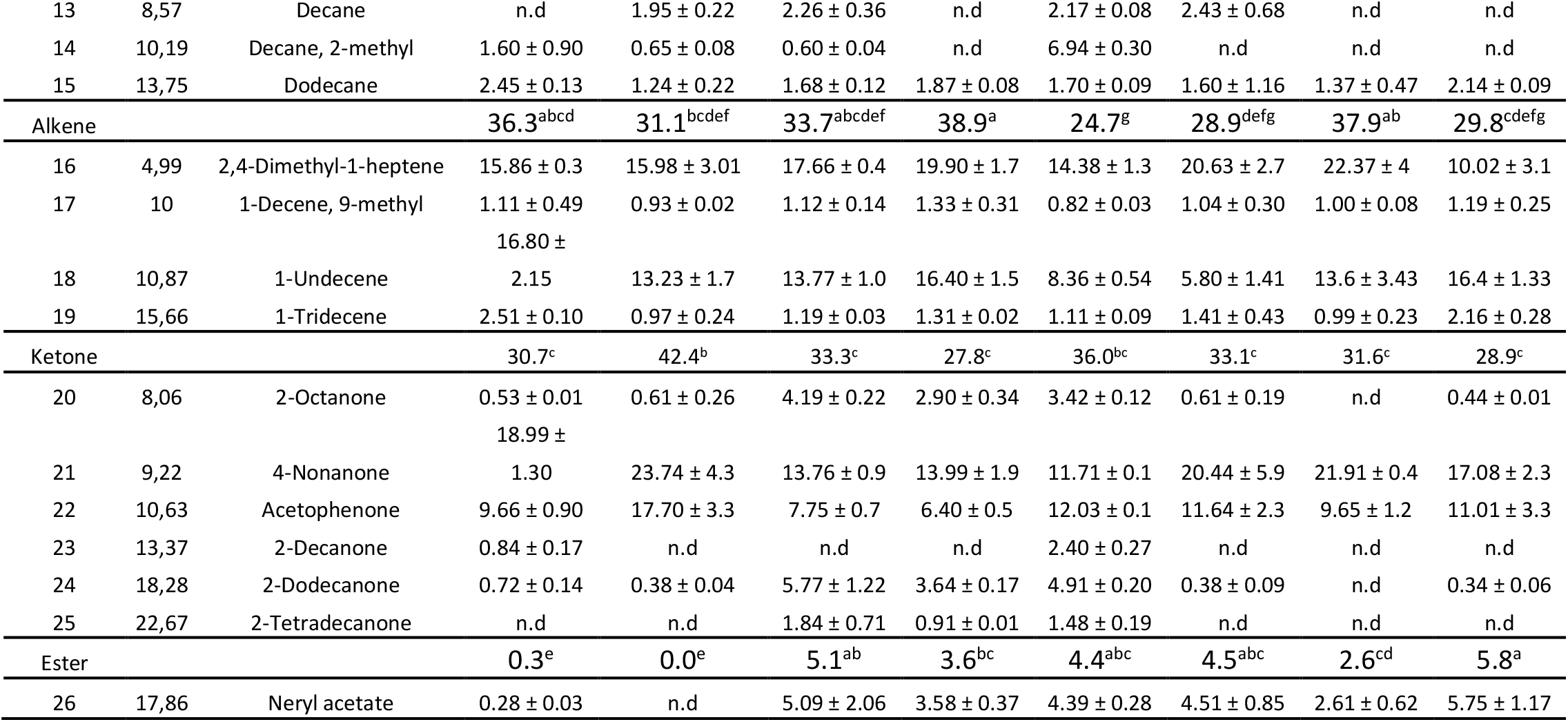
Identification and quantification as % area of VOCs from each CFS sample analysed. Results are expressed as mean ± standard deviation. Different letters indicate significant differences among treatments (P ≤ 0.01). The experiment was performed in triplicate (n=3).

Compared to the other samples, the control showed a different spectrum of compounds with the prevalence of an alkene (2,4 dimethyl-1-heptene and 1-undecanol) and a ketone (acetophenone). Fermentation of the MRS-B by the bacterial isolates determined an increase of the repertoire of compounds, including alcohols, aldehydes, esters and acids. In particular, the concentrations of acids and alcohols increased significantly, especially in CFSs of isolates identified as *L. plantarum* and *P. pentosaceus*. Interestingly, all CFSs produced decanal, a compound of the aldehyde class. It was detected in high proportion in CFSs of isolates FR6 and COR2 of *E. faecium* as well as in isolates OV16 and OV17 of *L. plantarum*. Differently from all the other classes of compounds, after the fermentation in all CFSs the percentage of ketones increased. The 4-nonanone was the ketone identified in greatest quantity. Control samples showed lower value of ketones, except for the acetophenone.

### 3.5 Chemometric analysis

To further characterize the isolates recovered from drupes, a PCA analysis based on VOC, organic and phenolic data of all CFSs was performed. The sum of all these components (PC) accounted for 45.8% of the total variance, while PC1 and PC2 represented 28.2% and 17.6% of the total variances, respectively. Figure 2 shows that the samples grouped according to their metabolic profile rather than the species as identified by peptide mass fingerprinting and sequencing of the 16S rRNA. The CFSs analysed were split into four different clusters, identified in relation to the relative abundance of a metabolite. The PC1 distributed the OV8, OV11 and COR 3 isolates on positive axis. These latter isolates on the loading plot (Fig. 2B) showed a cluster in relation to the production of benzonic acid (V 30), octane (V 20), 2-dodecanone (V 24), decane (V 13), undecanoic acid (V 3). Conversely, isolates of *L. plantarum* (OV 20, OV16, OV9) showed an aggregation type for the production. of 1-decene, 9-methyl (V 17), DL-3 phenyllatic acid (V 27) and 1,2-dihydroxybenzene (V 28).

**Figure 2.**
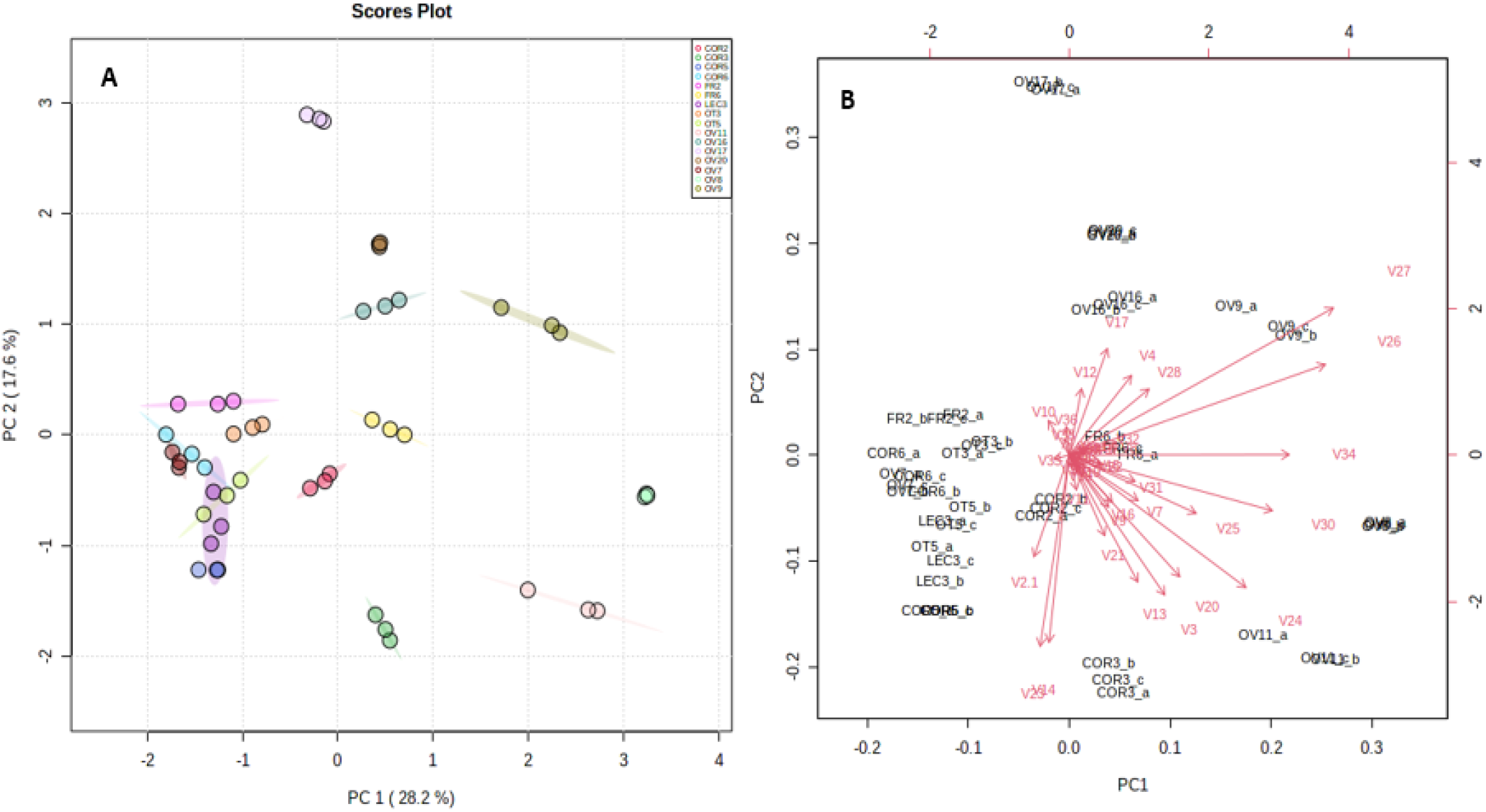
Principal Component Analysis (PCA), scores plot (a) and loading plot (b), based on volatile compounds, organic and phenolic acids of CFSs.

## 4. Discussion

In this study, it was demonstrated that fermentates of LABs isolated from olives exert a noticeable antifungal activity against a wide range of fungi *s*.*l*. pathogenic to olive or producing mycotoxins that may contaminate food products of olive industry, such as olive oil. They were effective against *A. alternata, A. flavus, C. acutatum, C. fiorinae, C. gloeosporioides, C. nymphaeae, F. oxysporum, P. digitatum, P. expansum, P. nordicum, Ph. nicotianae* and *Ph. oleae*. LABs isolated from olives in this study were identified as *L. plantarum, P. pentosaceus, E. faecium* and *S. salivarius*. There are several reports of the presence of *L. plantarum, P. pentosaceus* and *Enterococcus* sp. in different matrixes including table olive fruits (Benítez-Cabello et al., 2019; Heperkan, 2013; Hurtado et al., 2012). In particular, previously articles highlighted *L. plantarum* and *P. pentosaceus* showed antimicrobial activity (Dopazo et al., 2021; Passariello et al., 2020; Qi et al., 2021). However, all previous studies focused on LABs isolated from fruits or brine of olive table cultivars and their biotechnological applications in the food industry. By contrast, LABs characterised in this study were from olive oil varieties. Moreover, to the best of our knowledge, this is the first report of antifungal activity of *S. salivarum* and *E. faecium* and the first time that it is envisaged a potential application of LABs isolated from olive fruits and their fermentates in agriculture as potential BCAs or natural fungicides, respectively. As many of the fungi *s*.*l*. inhibited by LABs isolated from olive drupes, such as *A. alternata, C. acutatum, C. gloeosporioides, F. oxysporum* and *Ph. nicotianae*, are very polyphagous pathogens (Aloi et al., 2021; Biasi et al., 2016; Dean et al., 2012; Kamali-Sarvestani et al., 2022; Kamoun et al., 2015; Riolo et al., 2021) it can be hypothesized that their antifungal potential could be exploited to manage diseases of several other crop plants, besides olive. Chemical analyses of CFSs from liquid cultures of all LABs revealed they produce a large repertoire of active metabolites, including lactic acid, numerous phenolic acids and VOCs, that might be valued also in food industry with the same purposes for which the use of metabolites of other LABs is a consolidated technique. The antifungal activity of LABs recovered from olives varied considerably depending on the bacterial isolate and the fungi tested. For instance, no or a scarce antifungal activity was shown by all LABs tested against *Pl. tracheiphilus* and *A. flavus*. Most LAB isolates tested in this study inhibited significantly the *in vitro* growth of *Colletotrichum* species, but there were exceptions, such as those of C2 and RD9B isolates of *C. gloeosporioides* and *C. godetiae*. Similarly, *Penicillium* species showed a lower sensitivity than *Colletotrichum* species, but *P. nordicum* was an exception as its sensitivity to CFSs of most bacterial isolates was even lower than the sensitivity showed by *Colletotrichum* species. This sensitivity of *P. nordicum* to fermentates of LABs is consistent with the results of Guimarães et al. (2018). In this study, three distinct methods were used to assay the antifungal activity of LAB fermentates, the culture overlay, the agar diffusion and the serial dilution tests, the last one to determine both MIC and MFC values of CFSs. In general, results of the three tests showed a similar trend. However, also in this case there were exceptions. The most relevant one was the *E. faecium* strain FR6. In the culture overlay test this isolate showed no antifungal activity against *C. fioriniae, C. nympheae* and *F. oxysporum*, but showed a high inhibitory activity against these fungi in the agar diffusion and the serial dilution tests, suggesting the inhibitory activity was due to diffusible metabolites released by the bacterium in the culture medium. Some of the metabolites recovered from the culture liquids in this study have been previously reported as antifungal agents produced by LABs (Lavermicocca et al., 2003; Omedi et al., 2019; Wang et al., 2012). For instance, benzoic acid, cinnamic acid and its derivatives were reported in the literature to exert antifungal activity (Berne et al., 2015; Lima et al., 2018). Also caffeic acid, vanillic acid and vanillin were mentioned as phenolic acids with antifungal activity (Elansary et al., 2019; Fitzgerald et al., 2005; Li et al., 2022). The presence of hydroxycinnamic acids, p-coumaric, caffeic acid and ferulic acid is correlated to the capacity of *L. plantarum* to metabolize these phenolic compounds (Landete et al., 2021) while the lower level of acetophenone in CFSs, compared to the control samples, was probably correlated to the reduced potential of LABs to produce chiral alcohols (Baydaş et al., 2020). Interestingly, all CFSs produced decanal, a compound of the aldehyde class. Decanal was reported to possess antifungal activity against *P. digitatum, P. italicum* and fungi of other genera such as *Aspergillus* and *Geotrichum* (Li et al., 2022; Tao et al., 2014; Zhang et al., 2017; Zhou et al., 2014). Isolates of the genus *Lactiplantibacillus* characterised in this study produced a greater amount of lactic acid than LABs of other genera. Moreover, the chemometric analysis separated he CFSs of the LABs isolated from olives into four distinct clusters based on the relative abundance of specific metabolites. For instance, isolates OV8, OV11 (*L. plantarum)* and COR 3 *(P. pentosaceus)*, formed a cluster in relation to the production of benzonic acid, octane, 2-dodecanone, decane and undecanoic acid. Conversely, the isolates OV 20, OV16 and OV9 (as *L. plantarum)* grouped because of their ability to produce 1-decene, 9-methyl, DL-3 phenyllatic acid and 1,2-dihydroxybenzene. However, no obvious correlation was found between both the level and spectrum of antifungal activity showed by LAB isolates characterised in this study and their metabolic profiles.

The discovery that LABs isolated from drupes of olive oil varieties exert an effective inhibitory activity on a wide range of fungi *s*.*l*. is not only a novelty but open the way to potential biotechnological applications. Their characterization and the identification of active metabolites present in their fermentates are a preliminary step toward this objective.

