## Supplementary table S1 for "Antifungal activity of selected lactic acid bacteria from olive drupes"

| **Nº** | **Rt** | **Compound** | **Identification** | **LRI DB5** | **LRI Lit** | **Reference** |
| --- | --- | --- | --- | --- | --- | --- |
| **Acid** |  |  |  |  |  |  |
| 1 | 3,87 | Acetic acid | MS |  |  |  |
| 2 | 7,93 | Hexanoic acid | MS + LRI | 975 | 978 | Engel et al. (2007) |
| 3 | 19,66 | Undecanoic acid | MS + LRI | 1451 | 1460 | Pino et al. (2001) |
| **Alcohol** |  |  |  |  |  |  |
| 4 | 13,44 | 1-Nonanol | MS + LRI | 1187 | 1186 | Alissandrakis et al. (2007) |
| 5 | 15,34 | 1-Decanol | MS + LRI | 1264 | 1264 | Casarek et al. (2006) |
| 6 | 16,19 | 3-Undecanol | MS + LRI | 1299 | 1308 | Dickschat et al. (2004) |
| 7 | 18,02 | 1-Undecanol | MS + LRI | 1376 | 1376 | Jones et al. (2009) |
| **Aldehyde** |  |  |  |  |  |  |
| 8 | 14,01 | Decanal | MS + LRI | 1210 | 1208 | Alissandrakis et al. (2007) |
| **Alkane** |  |  |  |  |  |  |
| 9 | 3,54 | Heptane, 4-methyl | MS |  |  |  |
| 10 | 3,68 | Pentane, 2,3-dimethyl | MS |  |  |  |
| 11 | 4,42 | Heptane, 2,4-dimethyl | MS + LRI | 819 | 818 | Liu et al. (2006) |
| 12 | 7,57 | Nonane, 4-methyl | MS + LRI | 960 | 961 | Kotowska et al. (2012) |
| 13 | 8,57 | Decane | MS + LRI | 1000 | 1000 | Kohl et al. (2001) |
| 14 | 10,19 | Decane, 2-methyl | MS + LRI | 1062 | 1061 | Kotowska et al. (2012) |
| 15 | 13,75 | Dodecane | MS + LRI | 1200 | 1200 | Holldobler et al. (2004) |
| **Alkene** |  |  |  |  |  |  |
| 16 | 4,99 | 2,4-Dimethyl-1-heptene | MS + LRI | 848 | 843 | Goeminne et al. (2012) |
| 17 | 10,00 | 1-Decene, 9-methyl | MS + LRI | 1054 | 1055 | Zaikin & Borisov (2002) |
| 18 | 10,87 | 1-Undecene | MS + LRI | 1087 | 1090 | Song et al. (2003) |
| 19 | 15,66 | 1-Tridecene | MS + LRI | 1277 | 1283 | Luo et al. (2001) |
| **Ketone** |  |  |  |  |  |  |
| 20 | 8,06 | 2-Octanone | MS + LRI | 980 | 984 | Sampaio & Nogueia (2006) |
| 21 | 9,22 | 4-Nonanone | MS + LRI | 1025 | 1030 | Mahmood et al. (2004) |
| 22 | 10,63 | Acetophenone | MS + LRI | 1078 | 1078 | Buchin et al. (2002) |
| 23 | 13,37 | 2-Decanone | MS + LRI | 1184 | 1188 | Dhanda et al. (2003) |
| 24 | 18,28 | 2-Dodecanone | MS + LRI | 1388 | 1388 | Limberger et al. (2002) |
| 25 | 22,67 | 2-Tetradecanone | MS + LRI | 1594 | 1594 | Kurashov et al. (2014) |
| **Ester** |  |  |  |  |  |  |
| 26 | 17,86 | Neryl acetate | MS + LRI | 1369 | 1368 | Allegrone et al. (2006) |

**Table S1.** Assignment of GC–MS signals
